# A Penalized Linear Mixed Model with Generalized Method of Moments for Complex Phenotype Prediction

**DOI:** 10.1101/2021.10.11.463997

**Authors:** Xiaqiong Wang, Yalu Wen

## Abstract

Linear mixed models have long been the method of choice for risk prediction analysis on high-dimensional genomic data. However, it remains computationally challenging to simultaneously model a large amount of genetic variants that can be noise or have predictive effects of complex forms. In this work, we have developed a penalized linear mixed model with generalized method of moments (pLMMGMM) estimators for prediction analysis. pLMMGMM is built within the linear mixed model framework, where random effects are used to model the joint predictive effects from all genetic variants within a region. Fundamentally different from existing methods that usually focus on linear relationships and use empirical criteria for feature screening, pLMMGMM can jointly consider a large number of genetic regions and efficiently select those harboring variants with both linear and non-linear predictive effects. Through theoretical investigations, we have shown that our method has the selection consistency, estimation consistency and asymptotic normality. Through extensive simulations and the analysis of PET-imaging outcomes, we have demonstrated that pLMMGMM outperformed existing models and it can accurately detect regions that harbor risk factors with various forms of predictive effects.

## 1. Introduction

Accurate disease risk prediction plays an important role in precision medicine, an emerging model of healthcare that tailors treatment strategies based on individuals’ profiles (Ashley, 2015). Over the past decades, genome-wide association and wholegenome sequencing studies have detected many disease-associated genetic variants. While it is hoped that these genetic findings can facilitate the ongoing risk prediction studies (Speed and Balding, 2014; Weissbrod, Geiger and Rosset, 2016; Wen and Lu, 2020), existing genetic risk prediction models can only explain a small proportion of the heritability. The complex relationships between predictors and phenotypes (Speed and Balding, 2014; Weissbrod, Geiger and Rosset, 2016), the huge amount of noise in high-dimensional genetic data (Byrnes et al., 2013), and the high computational cost (Weissbrod, Geiger and Rosset, 2016; Wen and Lu, 2020) greatly limit the prediction accuracy of existing models.

Linear mixed models (LMMs) and their extensions are the most widely used methods for risk prediction studies (Speed and Balding, 2014; Weissbrod, Geiger and Rosset, 2016; Wen and Lu, 2020). The genomic best linear unbiased prediction (gBLUP) method, which was first introduced by Harris et al. (2008) to predict milk production and then extended for the prediction of human traits (Yang et al., 2010), is one of the earliest method within the LMM framework. gBLUP assumes that each genetic variant acts in an additive manner and their effect sizes follow the same normal distribution. gBLUP is equivalent to a LMM with one random effect term, in which the variance-covariance structure encodes the assumed linear additive relationships. gBLUP only needs to estimate one parameter associated with the random effect term, making it computationally efficient. While easy to implement, the modeling assumptions in gBLUP are too simple, and thus efforts have been made to relax them. For example, Speed and Balding (2014) have shown that genetic variants from different regions (e.g., eQTLs, intron SNPs and coding) can have different effect sizes, and thus extended the gBLUP to MultiBLUP, where the genome is split into multiple regions (e.g., based on gene or pathway annotations) with each being modeled by a random effect term with its own variance parameter. As converging evidences have suggested that epistasis widely exist (Buil et al., 2015; Moore and Williams, 2009), Weissbrod, Geiger and Rosset (2016) further generalizes the MultiBLUP via embedding cumulative predictive effects from each region into the reproducing kernel Hilbert space, where a pre-specified kernel function is used to construct variance-covariance matrices for each region, making it capable of capturing non-linear predictive effects. Recently, Wen and Lu (2020) incorporated the multi-kernel learnings into the LMMs, where complex predictive effects can be efficiently captured. While the advances in LMM-based methods offer greater flexibility in modeling complex diseases, their levels of successes are largely limited, mainly due to the high computational cost when a large number of random effects are used to accommodate different types of predictive effects.

While high-dimensional genomic data allow for thorough investigations of disease etiologies, they also contain a huge amount of noise. Within the LMM framework, using all genetic regions, including those that only harbor noise variants, to build risk prediction models not only substantially increases the computational complexity, but also reduces the robustness and accuracy of the models. Byrnes et al. (2013) has already demonstrated that in the absence of good biological annotations, variable selection is an efficient way in improving prediction accuracy, especially for high-dimensional data. Conventional variable selection methods (e.g., BIC, GIC, and forward selection) as well as those empirical criteria employed in LMMs can all be used to select predictive regions (Speed and Balding, 2014; Weissbrod, Geiger and Rosset, 2016). However, there is no theoretical guarantee for their optimal performance. *L*_1_ penalization is a common technique used for simultaneously selecting predictive variables and estimating their effect sizes (Ghosh and Chinnaiyan, 2005; Ma, Song and Huang, 2007; Wu et al., 2009; Sun and Wang, 2012). Recently, it is introduced to LMM-based risk prediction models, where penalties are imposed on the random effect terms to allow for consistent and efficient selection of predictive regions (i.e., random effects) (Wen and Lu, 2020; Li, Lu and Wen, 2020). While these advances can reduce the impact of noise and improve prediction accuracy, their parameter estimations can be computationally demanding. Existing algorithms usually adopt the one-step approximation procedure (Fan and Li, 2001; Zou and Li, 2008) and their performance depends on the initial values that are usually set to be either the maximum likelihood estimators (MLE) or the restricted maximum likelihood estimators (REML). However, both MLE and REML can itself be hard to obtain for LMMs, when a large number of random effects is considered. As a consequence, existing penalized LMMs can only handle a few regions with limited types of predictive effects, leading to sub-optimal prediction performance.

The parameters in LMM-based models are usually estimated with either MLE or REML (VanRaden, 2008; Yang et al., 2010; Speed and Balding, 2014), where Newton-Raphson or expectation-maximization algorithms are commonly used to optimize the objective functions. Although both MLE and REML are statistically efficient, their estimation procedure involves repeatedly inversing the variance-covariance matrix, making it computationally prohibited to consider a large amount of random effects. Recently, simulated annealing algorithms have also been introduced to optimize the objective function of LMMs (Weissbrod, Geiger and Rosset, 2016). However, the performance of simulated annealing algorithms is determined by empirical criteria, and thus it will be very likely to find a local optimal or it will last for a very long time to find a global optimal. Generalized method of moments (GMM) are long existing alternatives to REML/MLE for LMMs (Rao, 1970, 1971, 1972), where statistical efficiency is traded with computational efficiency. It is a promising alternative for penalized LMMs with multiple random effects, as it can change the objective function into a quadratic form that is much easier to optimize. Indeed, GMM estimators have been used in LMMs for the estimation of variance components. For example, Zhu and Weir (1996) used the minimum norm quadratic unbiased estimation (MINQUE) method to estimate variance components for maternal and paternal effects in a bio-model for diallel crosses. Zhou (2017) unified the method of moments, the MINQUE criterion, the Haseman-Elston regression and the LD score regression to estimate the heritability of height using the Australian GWAS data. Pazokitoroudi et al. (2019) presented a randomized multi-component version of the classical Haseman-Elston regression, which uses method-of-moments to estimate heritability of 22 complex traits using the UK Biobank data. While GMMs have demonstrated their computational efficiencies in variance component estimation, they have not been used for the parameter estimations in penalized LMMs with multiple random effects, and their theoretical properties are rarely studied.

To address these issues, we developed a penalized LMM with GMM estimators (referred as pLMMGMM) to simultaneously select predictors and estimate their effect sizes in the prediction analysis. Similar to existing LMMs, the proposed method splits the genome into multiple regions and models the cumulative predictive effects for each region via random effect terms. Fundamentally different from existing LMMs that rely on MLE or REML, our method estimates its parameters using GMM, making it much more computationally efficient. Therefore, our method can 1) use a data-driven approach to choose appropriate kernel functions to reflect different types of relationships between predictors and outcomes, and 2) simultaneously and efficiently model a large number of regions (i.e., random effects) and detect those that are predictive. In the following sections, we first presented the pLMMGMM model and its theoretical properties in section 2. In section 3, we conducted extensive simulation studies to evaluate its empirical performance, and further compared its prediction accuracy with commonly used methods. In section 4, we illustrated the practical utility of our method by analyzing a dataset obtained from Alzheimer’s Disease Neuroimaging Initiative (ADNI) study (Saykin et al., 2010).

## 2. Methods

LMMs have long been used for risk prediction analysis on high-dimensional genomic data (de los Campos et al., 2013; Speed and Balding, 2014; Weissbrod, Geiger and Rosset, 2016). For completeness, we first presented the LMMs used for prediction research, and then introduced our penalized LMM where the parameters are estimated using GMM. Finally, we showed the theoretical properties of our proposed GMM-based estimators.

### 2.1. Linear Mixed Model for Risk Prediction with Genomic Data

The fundamental assumption in LMMs is that genetically similar individuals can have similar phenotypes. As genetic variants located at different locations (e.g., eQTLs, intron SNPs and coding) can have different effect sizes (Speed and Balding, 2014), we first split the genome into *R* regions based on some criteria (e.g., gene annotation and pathway), and model the outcomes ***Y*** as:

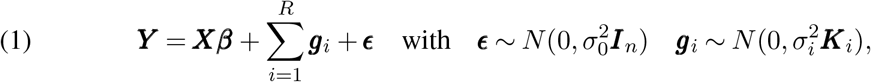

where ***X*** is an *n* × *P* matrix of the demographic variables (e.g., age and gender), and ***β*** is their effect sizes. ***g***_*i*_ is the cumulative predictive effect from the *i*th region, and ***K***_*i*_ is a *n* × *n* kernel matrix measuring the genetic similarities of region *i*.

Similar to existing LMMs, the variance-covariance matrix for each cumulative effect implicitly determines the assumed relationship between predictors and the outcome. For example, if a linear kernel is used for each region (i.e., 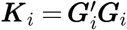 with ***G***_*i*_ is an *n* × *p_i_* genotype matrix for region *i*), the proposed model in equation 1 is equivalent to

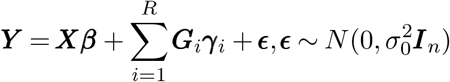

where 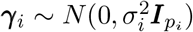. Therefore, a linear kernel implicitly assumes that there is a linear additive relationships between predictors and the outcome. To accommodate more complex relationships, we extended the single kernel into multiple kernels, where multiple kernel matrices are combined in a data-driven manner. For example, when both linear and pair-wise interaction effects are considered, we set the variance-covariance matrix to be 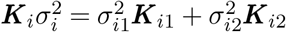, where ***K***_*i*1_ is a linear kernel designed to capture additive effects and **K**_i2_ is the polynomial kernel with 2 degree of freedom designed to model the pairwise interaction effects. More kernels (e.g., the RBF kernels) can be added to the candidate kernel set to efficiently model various types of predictive effects. By using random effects to capture cumulative predictive effects and kernelizing the covariance matrices, our proposed method not only reduces the model parameters, but also offers a very flexible framework for modeling traits with various underlying genetic architecture (Wen and Lu, 2020; Li, Lu and Wen, 2020).

### 2.2. Penalized Linear Mixed Model with Generalized Method of Moments Estimators

Converging evidence has shown that not all genetic variants and regions are predictive (Wen and Lu, 2020; Wen et al., 2016; Li, Lu and Wen, 2020; Speed and Balding, 2014; Weissbrod, Geiger and Rosset, 2016). Including noise can reduce the robustness and accuracy of the prediction model. For our proposed model, if region *i* is not predictive, then there is no variations for their cumulative predictive effect ***g***_*i*_ and thus 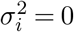. Therefore, selecting predictive regions is equivalent to determine which 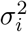 is not zero.

*L*_1_ penalty is a commonly used technique for simultaneously selecting predictors and estimating their effect sizes. For example, Wen and Lu (2020) added an *L*_1_ penalty to the log likelihood function of LMMs to simultaneously select predictive regions and estimate their effect sizes:

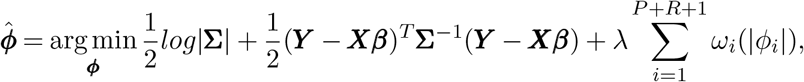

where 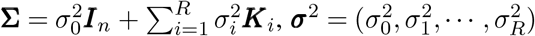, and ***ϕ*** = (***σ***^2^, ***β***). While capable of detecting predictive regions, this method can only consider limited number of genetic regions in practice, mainly due to their high computational cost.

To simultaneously model predictive effects from a large number of genetic regions (i.e., random effects in LMMs), we proposed to use the generalized method of moments to select predictive regions and estimate their effect sizes. Clearly, the variance of ***Y*** depends on covariates ***X***, and thus we propose to follow the same procedure developed by Pazokitoroudi et al. (2019) to choose an ***A*** matrix, such that the variance of ***A’Y*** is independent of the covariates ***X***. Let ***V*** = ***I*** – ***X*** (***X’X***)^−1^***X’*** is a symmetric and idempotent of rank *n* – *P* matrix. We consider the eigen decomposition of ***V*** = ***EDE’***, where ***D*** is a diagonal matrix with *n* – *P* ones and *P* zeros on the diagonal. We set the ***A*** matrix as the first *n* – *P* columns of ***E***. Therefore, ***AA’*** = ***V***, ***A’A*** = ***I***, ***A’X*** = 0, and

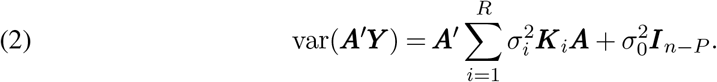

Traditionally, the parameters in equation 2 are estimated using REML estimators that can be computationally infeasible when the number of regions is large. To overcome the computational bottleneck, we propose to trade statistical efficiency with computational efficiency, and develop a penalized GMM estimator for its parameter estimation:

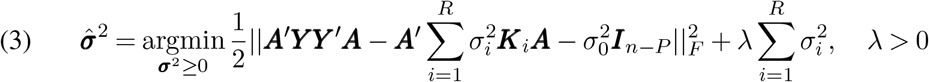

By using the proposed penalized GMM estimators, the objective function is in a quadratic form that is much easier to optimize as compared to the traditional REML or MLE estimators. It is straightforward to see that the parameters can be easily obtained via solving equation 4 (the details are shown in appendix A.1):

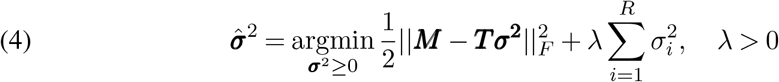

where ***M*** = *vec*(***A’YY’A***), ***T***_*i*_ = *vec*(***A’K_i_A***), ***T***_*i*_ is the *i*th column of ***T*** matrix, and thus ***T*** = (***T***_0_,***T***_1_,…,***T***_*R*_). *vec*(.) is the vectorization of a matrix. Equation 4 can be solved by the coordinate descent algorithm implemented in `glmnet` R package (Friedman, Hastie and Tibshirani, 2010).

Let ***Y***_*a*_ = (***Y***_*p*_,***Y***), where ***Y*** is the *n* × 1 vector of outcomes in the training data and ***Y***_*p*_ is *n_p_* × 1 vector of outcomes to be predicted. Given the parameter estimates for ***σ***^2^ and ***β***, the variance of ***Y***_*a*_ can be directly derived as 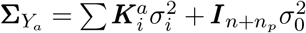, where 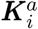 is the (*n_p_* + *n*) × (*n_p_* + *n*) genetic similarity matrix calculated from all samples. The variance of ***Y***_*a*_ can be written as,

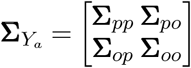

where **Σ**_*pp*_ and Σ_*oo*_ are the variance matrices for the testing and training samples, respectively. **Σ**_*po*_ is the covariance matrix between testing and training samples. Therefore, the predictive values for the testing samples can be calculated as,

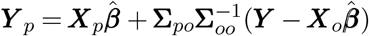

where the ***X***_*p*_ and ***X***_*o*_ are the covariates of testing samples and training samples, respectively.

### 2.3. Theoretical Properties

We investigated the theoretical properties of our proposed method, including the selection consistency, estimation consistency and asymptotic normality. We will explore whether our model can choose the right predictive variables, and establish the asymptotic distribution of our estimators.

#### 2.3.1. Notations and Assumptions

Let 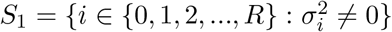 denote the set of all predictive regions, and 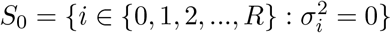 be the set of regions that just harbor noise variants. Let 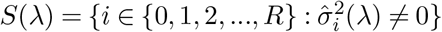 be the estimated set of predictive regions for a given value of λ that is selected independent of predictors and outcomes. For simplicity and without loss of generality, we assume the first *q* regions are predictive and the remaining *R* – *q* regions are noise. Let ***T***(1) = (***T***_0_,***T***_1_,…,***T***_*q*_) and ***T***(2) = (***T***_*q*+1_, ***T***_*q*+2_,…, ***T***_*R*_). Let 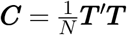, where *N* is the number of rows in ***T*** matrix. Therefore, ***C*** can be written as

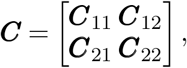

where 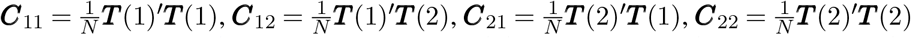.

Similar to Wu, Yang and Liu (2014), we assumed the following conditions for our method:

#### Assumption 1.

*There exist constants 0 ≤ c_1_ ≤ c_2_ ≤ 1, 0 ≤ c_3_ < c_2_ – c_1_, and M_1_, M_2_, M_3_ > 0, such that the following conditions hold*,

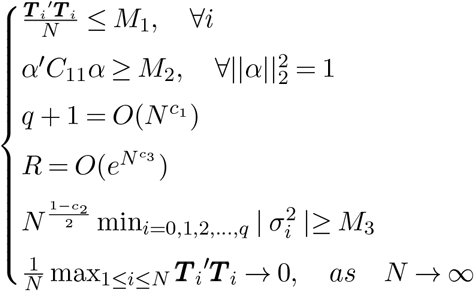

##### Assumption 2.

*Nonnegative irrepresentable condition: there exists a positive constant vector **ρ**, such that*

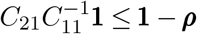

##### Assumption 3.

*Restricted eigenvalue condition: there exists constant k_m_, such that:*

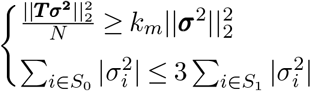

##### Assumption 4.

*Column normalization condition:*

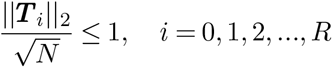

#### 2.3.2. Theoretical Properties

##### Theorem 2.1

(Variable selection consistency). *Under the assumptions 1 and 2, the pLMMGMM method can have variable selection consistency. In particular, when* 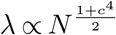 *where c_3_ < c_4_ < c_2_ – c_1_, the following condition holds:*

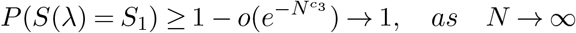

The theorem 2.1 demonstrates that the proposed pLMMGMM can consistently select the predictive regions even when the number of regions (i.e., *R*) increases faster than the sample size (*N*) at an exponential speed. The proof of the selection consistency can be seen in the appendix A.2.

##### Theorem 2.2

(Variable estimation consistency). *Under the assumptions 3 and 4, the pLMMGMM method can have variable estimation consistency. In particular, when the regularization parameter* 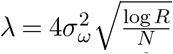, *there exists constants υ*_1_, *υ*_2_ > 0 *such that, with probability at least 1 – υ*_1_ exp(−*υ*_2_*N*λ^2^),

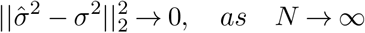

Theorem 2.2 indicates that the penalized GMM estimators has variable estimation consistency. The details of the proof for theorem 2.2 can be seen in the appendix A.3.

##### Theorem 2.3

(Asymptotic normality). *Under the same setting in theorem 2.1, the nonzero estimator* 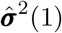 *is asymptotically normal*,

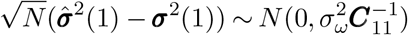

Theorem 2.3 indicates that the penalized GMM estimators for those nonzero parameters are asymptotically normally distributed. The proof for theorem 2.3 can be seen in the appendix A.4.

## 3. Simulation Studies

We investigated the performance of the proposed pLMMGMM through extensive simulation studies, where the impacts of noise and the underlying disease models are evaluated. We considered sample sizes of 500 and 1000. For each setting, we randomly chose 70% of the samples to train the model, and used the remaining to assess the prediction accuracy, which was measured by both Pearson correlations and mean square errors (MSE). We further compared our method with two widely used methods, including gBLUP with its default settings (Yang et al., 2010) and MKLMM (Weissbrod, Geiger and Rosset, 2016). Note that by default, MKLMM only includes a pre-determined number of regions that are selected based on the rank of LR-ratio for each region. Therefore, for a fair comparison, we compared our method with MKLMM under two settings, where the screening step is included (i.e., the default of MKLMM) or omitted (i.e., all regions are considered jointly). We denoted these two settings as MKLMM and MKLMMpre, respectively. We did not compare our method with MultiBLUP, mainly because MultiBLUP is equivalent to MKLMM with a linear kernel. To evaluate whether the proposed method can select predictive regions, we calculated the sensitivity and specificity for our method. For all simulations, to mimic the real human genome, we directly obtained genotypes from the 1000 Genome Project (The 1000 Genomes Project Consortium, 2015), and constructed each region with 30 randomly selected SNps that are within 75Kb.

### 3.1. Scenario I: The Impact of The Number of Noise Regions

In this set of simulations, we gradually increased the number of noise regions to evaluate their impact. In particular, we randomly set two regions as causal, and simulated the outcomes under an additive model:

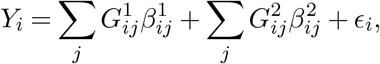

where 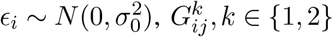 represents the *j*th SNPs on the *k*th causal region for individual *i*, and 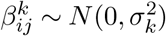 is their corresponding effect. It is straightforward to show that

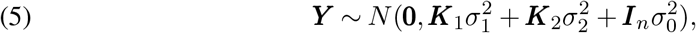

where 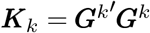 and ***G**_k_* is the genotype matrix for region *k*. Therefore, as shown in equation 5, we simulated the outcomes based on a multivariate normal distribution.

We gradually increased the number of noise regions from 0 to 98 (i.e., the total number of regions ranges from 2 to 100). For each setting, we conducted 1000 Monte Carlo simulations, and calculated the prediction accuracy based on testing samples. We reported the Pearson correlation, MSE, and the computational cost for each method. We further calculated the probabilities of correctly detecting causal and noise regions for our method.

The Pearson correlations and MSEs for sample sizes of 500 and 1000 are shown in Figure 1 and Supplementary Figure S1, respectively. Among all the scenarios considered, pLMMGMM performs the best. When there is no noise regions, our proposed method has similar levels of Pearson correlations as those of gBLUP and MKLMM, but its MSEs tend to be smaller. As the number of noise regions increases, the prediction accuracy of the proposed method remains roughly stable, while the performance of the other methods decreases to some extent with gBLUP being affected the most and MKLMM being the least. gBLUP assumes that all genetic variants act in additive manner and their effect sizes follow the same normal distribution. Therefore, as the number of noise regions increases, the assumption of gBLUP has been severely violated, and thus its performance dropped the most. On contrary, MKLMM allows genetic variants from different regions having different effects, and thus its variance component estimates in LMM can differ substantially for variants located on different regions. Therefore, it has certain capacities in dealing with noise regions, and it tends to have better performance than gBLUP. Comparing MKLMM under the two settings, the default has better performance than the one where the screening step is omitted. The employed pre-screening procedure can limit the number of regions, and thus reduces the number of random effects in LMM, which improves the robustness and computational efficiency of MKLMM. However, the screening step implemented in MKLMM considers one region at a time and relies on some empirical criteria for region pre-selection, both of which may lead to a sub-optimal prediction model.

**Fig 1.**
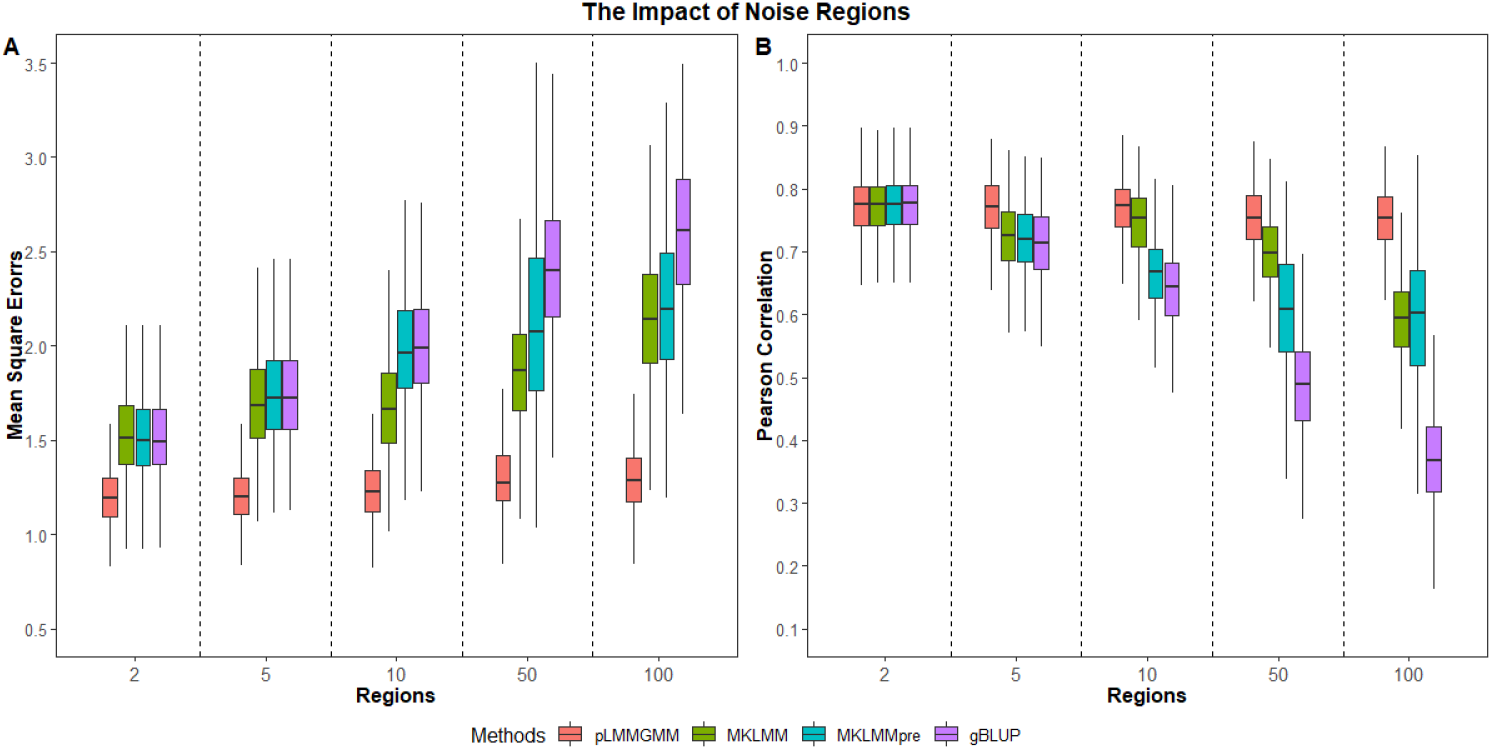
The impact of the number of noise regions on Pearson correlations and MSEs (n = 500).

Comparing pLMMGMM with both gBLUP and MKLMM, our method can jointly consider a large number of regions and select those that are predictive. Therefore, its performance is relatively robust against the noise. Indeed, excluding noise regions cannot only improve prediction accuracy, but also improve the robustness of the model. With regards to variable selection, our pLMMGMM method can not only correctly detect those causal regions, but also identify those that are noise. The results for variable selection under 500 samples and 1000 samples are summarized in Table 1 and Supplementary Table S1, respectively.

**Table 1.** The chances of selecting two predictive regions as the number of noise regions increases (n = 500).

| Regions | Sensitivity | Specificity |
| --- | --- | --- |
| 5 | 1.000 | 0.905 |
| 10 | 0.999 | 0.884 |
| 50 | 1.000 | 0.909 |
| 100 | 0.999 | 0.932 |

One of the benefits of using GMM is to improve the computational efficiencies, and thus we compared the running time of our method and MKLMMpre, where the same LMM model was fitted using the REML estimator. While the Newton-Raphson algorithm is the most widely used method for optimizing the objective function of REML estimators, it can barely converge when the number of random effects is large. Indeed, the convergence rates for 2, 5, 10, 50, and 100 regions (i.e., random effects) are 100%, 15.5%, 7.9%, 0% and 0%, respectively. For REML estimated using the simulated annealing algorithm, it can converge. However, there is no guarantee for the simulated anneal algorithm to get the global optimum. As shown in Figure 1, MKLMMpre has lower prediction accuracy than our proposed method. In addition, the computational time of REML grows much faster than the GMM-based estimators as the number of random effects increases, regardless of the sample sizes considered (Figure 2 and Supplementary Figure S2).

**Fig 2.**
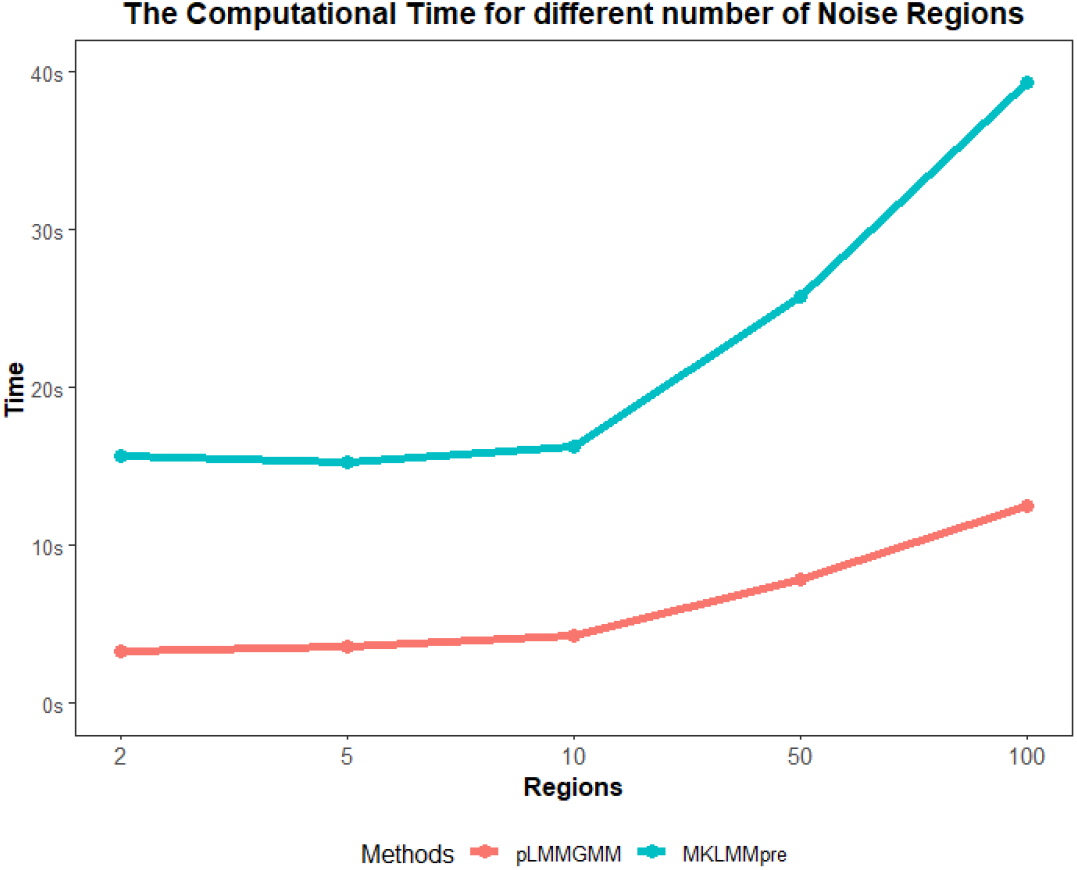
The impact of the number of noise genomic regions on computational time (n = 500).

### 3.2. Scenario II: The Impact of Disease Models

Complex traits and diseases are affected by a large number of genes through complicated biological pathways that are usually unknown in advance (Chatterjee et al., 2013). It has long been recognized that a risk prediction model with flexible modeling assumptions is more robust and accurate across a range of phenotypes with different genetic architectures. In this set of simulations, we evaluated the performance of our proposed method given different disease models. Similar to session 3.1, we considered 2 causal genes and simulated the outcomes using equation 5, where the kernel matrices ***K***_1_ and ***K***_2_ were used to reflect different disease models. Specifically, we considered 5 disease models, including 1)*L* + *L*: genetic variants on both regions have linear additive effects (i.e., *k_l_*(***x***_1_,***x***_2_) =<***x***_1_,***x***_2_>); 2) *R* + *R*: predictors from both regions have non-linear predictive effects (i.e., 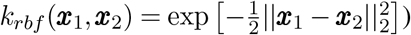; 3) *P* + *P*: both regions harbor variants with pair-wise interaction effects (i.e., *k_p_*(***x***_1_,***x***_2_) = (< ***x***_1_,***x***_2_ >)^2^); 4) *L* + *R*: genetic variants on the first and second regions have linear additive effects and nonlinear effects, respectively; and 5) *L* + *P*: predictors on the first and second regions have linear additive and pair-wise interaction effects, respectively. The details for each model setting and the corresponding kernels are summarized in Supplementary Table S2. In addition to these 2 causal regions, we also simulated 48 noise regions (i.e, the total number of regions were 50) for this set of simulations.

For MKLMM, different from the first simulation where only linear kernel is used, we used the default setting where the most appropriate kernels are selected in a data-driven manner. Note that since MKLMMpre performs worse than MKLMM that adopts an empirical screening rule (Figure 1 and Supplementary Figure S1), we only compared our method to MKLMM with screening implemented. Similar to simulation 1, we conducted 1000 Monte Carlo simulations for each setting, and summarized the prediction accuracy for all methods considered. In addition, for our proposed method, we also calculated the probabilities of detecting causal and non-causal regions.

As shown in Figure 3 and Supplementary Figure S3, the proposed pLMMGMM has the lowest MSEs and highest Pearson correlations among all methods considered, which indicates that our method is quite robust against different disease models. This is mainly because that our method can not only select predictive genetic regions, but also account for non-linear effects through selecting appropriate kernel functions from the candidate kernel set.

**Fig 3.**
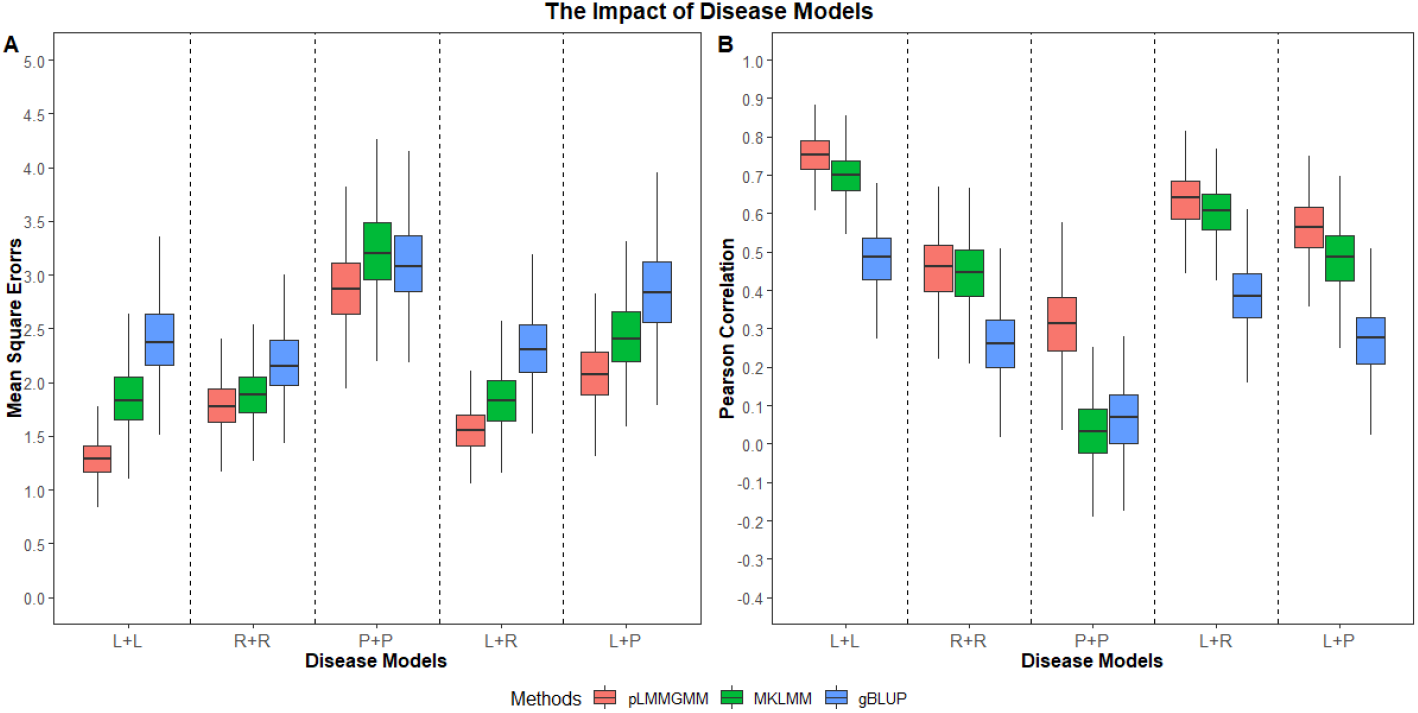
The impact of disease models. **L+L:** genetic variants on both regions have linear additive effects. **R+R:** predictors from both regions have non-linear predictive effects. **P+P:** both regions harbor variants with pairwise interaction effects. **L+R:** genetic variants on the first and second regions have linear additive and non-linear effects, respectively. **L+P:** predictors on the first and second regions have linear additive and pair-wise interaction effects, respectively (n = 500).

While MKLMM is designed to capture non-linear effects, in practice it can barely capture them. Indeed, for most of the non-linear models that we considered, the chance of selecting only linear kernels by the adaptive MKLMM is extremely high (> 99%). For example, for the disease model that has pair-wise interaction effects (i.e., *P* + *P*) on both causal regions, the chance of selecting the ideal polynomial kernel for MKLMM is close to 0%, whereas our proposed method has an average of 76% of choosing the appropriate polynomial kernel. As a consequence, the adaptive MKLMM performs very similar to a MKLMM that only considers the linear kernel, and it can perform worse than the MKLMM with the most appropriate kernel employed. See supplementary Figures S4 and S5, where the most appropriate kernels are used for MKLMM (i.e., the kernels are pre-determined based on the underlying etiology).

One of the key features of MKLMM is that it screens the genome and selects those that are predictive to build the risk prediction model. However, its ability in detecting causal regions also depends on the underlying disease model. For example, for the *P* + *P* disease model, the chance for adaptive MKLMM to select any of those two causal regions is very low (i.e., only noise regions are used for prediction), leading to a prediction model that performs even worse than gBLUP. For other disease models, although MKLMM is unlikely to efficiently capture those non-linear effects (i.e., not able to choose the most appropriate kernel), it has a relatively high chance of detecting the causal regions, and thus reduces the impact of noise. As a result, the adaptive MKLMM can outperform gBLUP under these settings, as gBLUP utilizes all regions and ignores the impact of noise. To evaluate whether our method can detect causal regions under different underlying disease models, we calculated the sensitivity and specificity, and the results are summarized in Table 2 and Supplementary Table S3. On average, our method achieves a sensitivity of 93% and a specificity of 95% among all the models considered under 500 samples, and a sensitivity of 99% and a specificity of 93% among all the models considered under 1000 samples.

**Table 2.** The chances of selecting two predictive regions under different disease models (n = 500).

| Disease models | Sensitivity | Specificity |
| --- | --- | --- |
| $S_1 : L + L$ | 0.999 | 0.919 |
| $S_2 : R + R$ | 0.969 | 0.951 |
| $S_3 : P + P$ | 0.794 | 0.980 |
| $S_4 : L + R$ | 0.959 | 0.931 |
| $S_5 : L + P$ | 0.917 | 0.949 |

## 4. Real Data Application

We used our proposed pLMMGMM method to predict positron emission tomography imaging outcomes, including FDG and AV45, using the whole-genome sequencing data obtained from the ADNI. ADNI is a longitudinal study designed for the prevention and treatment of Alzheimer’s Disease (AD) (Mueller et al., 2005). It measures clinical, imaging, genetic and biochemical biomarkers from each participant to investigate the pathology of AD. DNA samples from 818 participants aged between 55 and 90 were collected and sequenced on the Illumina HiSeq2000 at a non-Clinical Laboratory Improvements Amendments (non-CLIA) laboratory (Saykin et al., 2015). Genetic variants with missing rate large than 1% were first excluded, and then the remaining variants were annotated based on GRch37 assembly. We selected 95 AD-related genes based on existing literature (details are listed in Supplementary Table S4), and a total of 117,668 variants were included in the final analyses.

For our analyses, we are interested in predicting PET-imaging outcomes, including FDG and AV45, using the whole-genome sequencing data. We removed individuals that are either correlated or have missing outcomes, and the distributions of FDG and AV45 for the remaining samples (*n* = 539 for FDG and *n* = 401 for AV45) are shown in Supplementary Figure S6. To evaluate the prediction accuracy, we randomly split the samples into testing (*n* = 100) and training sets, where the training samples were used to train the prediction model and the remaining was used to calculate the Pearson correlations and MSEs. To avoid chance finding, we repeated this process 100 times.

The prediction accuracies for FDG and AV45 are shown in Figure 4. For both FDG and AV45, pLMMGMM has lower MSEs and higher Pearson correlations than both gBLUP and MKLMM. This indicates simultaneously considering multiple genes and excluding those that are not predictive can improve the robustness and accuracy of the risk prediction model.

**Fig 4.**
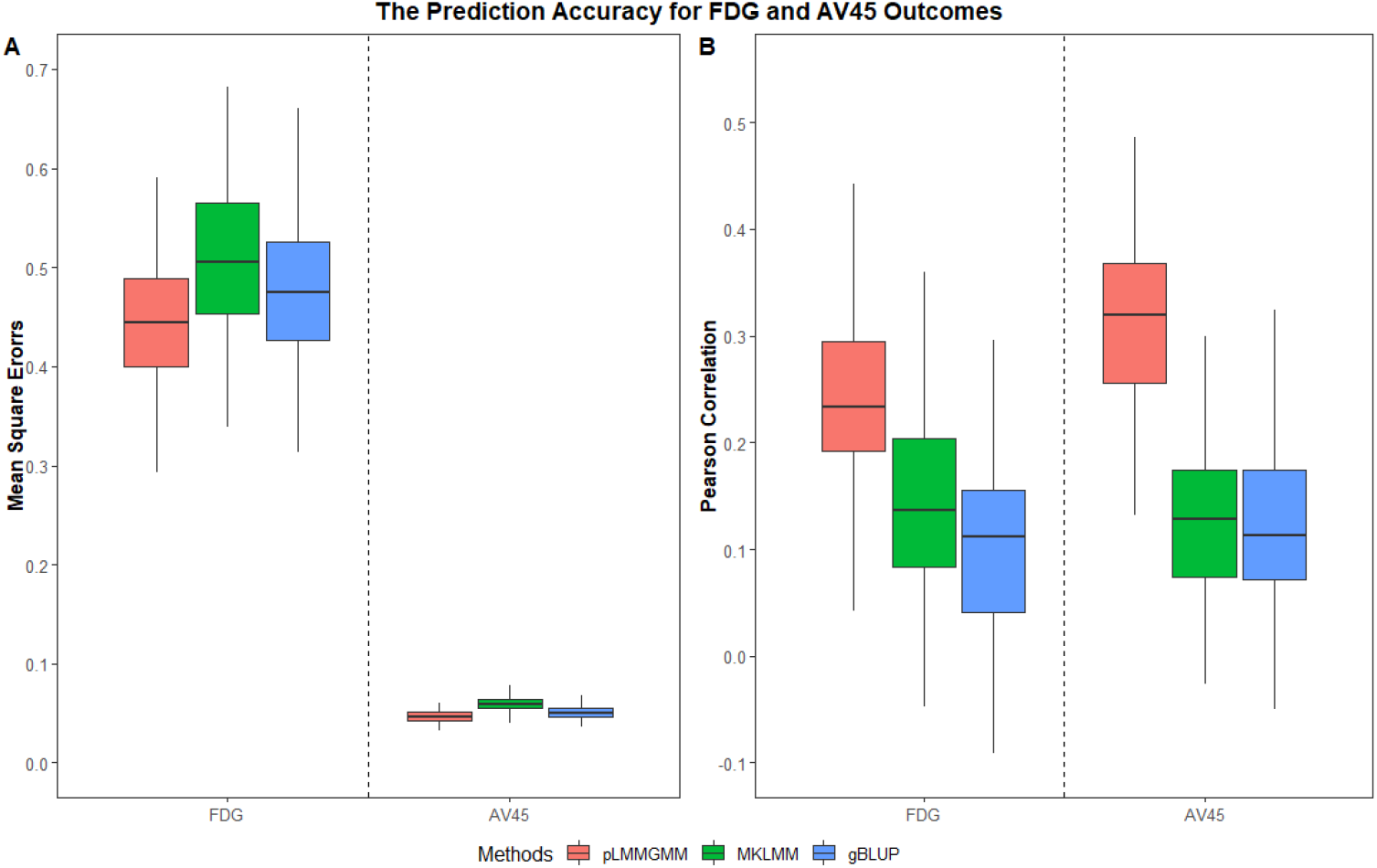
Accuracy comparison for FDG andAV45 outcomes.

Table 3 listed three genes that are most highly selected by pLMMGMM for either AV45 or FDG, and the selection details of all the genes are shown in Supplementary Table S4. For FDG outcomes, both *APOC1* and *APOE* are selected 100% times, and the remaining genes are averagely selected less than 1%. For AV45, *APOC1, APOE* and *TOMM40* are selected more than 97% times and the remaining genes are selected less than 5%. The highly selected genes, *APOC1, APOE* and *TOMM40*, are well-known AD-related genes (Ossenkoppele et al., 2013; Roses, 2010). All of the three genes are located on chromosome 19 and they have been widely known as genetic risk factors of AD. For example, van Duijn et al. (1994) found that *APOEϵ4* is highly associated with a group of 175 early-onset AD patients. Zhou et al. (2014) found the association of *rs*11568822 on *APOC1* gene with the increased AD risk in Caucasians, Asians and Caribbean Hispanics. Huang et al. (2016) found that *rs*2075650 on *TOMM40* is associated with AD patients for Caucasian and Asian subjects.

**Table 3.**
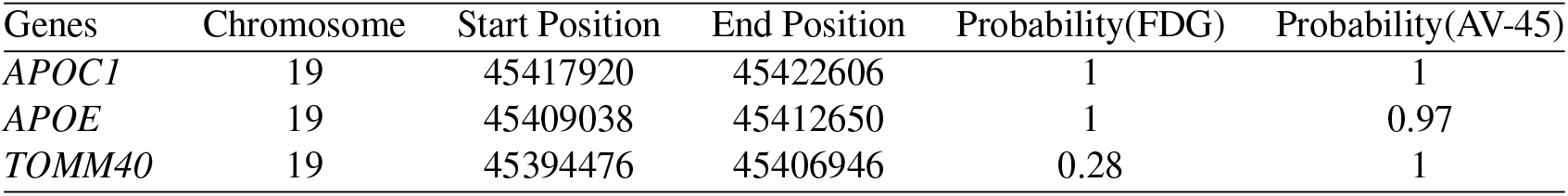
The top 3 genes highly selected for FDG and AV45 outcomes

## 5. Discussion

In this work, we presented a novel and computationally efficient penalized linear mixed model with generalized method of moments estimators for the prediction modeling on high-dimensional genomic data. The proposed pLMMGMM first splits the genome into multiple regions and then adopts multiple kernels for each region to capture complex predictive effects. pLMMGMM simultaneously models the joint predictive effects from all variants within each region, and efficiently select those that are predictive via a GMM-based estimator. Through theoretical proof, we have shown that our proposed method can achieve the consistency of variable selection and variable estimation. We also show that our proposed GMM estimators are asymptotically normal. Through extensive simulation studies and the analysis of ADNI dataset, we have demonstrated that our method 1) is more accurate and robust against various underlying disease models; 2) can accurately detect predictive regions; and 3) is much more computationally efficient, especially when the number of regions is large (i.e., the number of random effects is large).

The genomic data is high-dimensional and contains a large amount of noise. Including noise variants in the analyses can reduce the robustness and accuracy of a risk prediction model (Byrnes et al., 2013). Within the LMM framework, gBLUP cannot perform variable selection (Yang et al., 2010), and it subjects to low prediction accuracy when a large amount of noise present. Other LMM-based methods (e.g., MultiBLUP and MKLMM) select regions based on empirical criteria (Speed and Balding, 2014; Weissbrod, Geiger and Rosset, 2016), which cannot guarantee the optimal prediction performance. Existing penalized LMMs can detect predictive regions and estimate their effects, but they can only handle very limited number of regions (Wen and Lu, 2020; Li, Lu and Wen, 2020). On contrary, our proposed model can handle a large number of regions and efficiently remove those that are not predictive. We have proved that the probability to correctly identify all predictive regions approaches to 1, when the sample size is large. In addition, the effect estimates for these predictive regions are unbiased and asymptotically normal. As shown in the first simulation (Figure 1 and Supplementary Figure S1), the prediction accuracy for pLMMGMM remains stable as the amount of noise increases, whereas it can be greatly affected for other methods (i.e., gBLUP and MKLMM). Furthermore, the proposed pLMMGMM has achieved relatively high sensitivity and specificity, regardless of the number of noise regions (Table 1 and Supplementary Table S1).

The underlying genetic etiology for many human diseases is complicated and is usually unknown in advance. While it is widely accepted that models with flexible modeling assumptions can achieve more robust and accurate prediction performance across a range of phenotypes (VanRaden, 2008; Yang et al., 2010; Speed and Balding, 2014), existing LMMs mainly focus on linear relationships and thus their performance can be sub-optimal when non-linear predictive effects are present. The recent development in LMMs aims at capturing these nonlinear predictive effects via embedding them into the reproducing kernel Hilbert space, where appropriate kernel functions that reflect the underlying disease etiology are used (Weissbrod, Geiger and Rosset, 2016). However, how to pre-choose appropriate kernel functions can be challenging in practice, as they can be quite disease/trait dependent. In this work, we adopted the idea used in multi-kernel learning algorithms and put multiple kernels into a candidate set, from which appropriate kernel(s) are chosen through a data-driven approach. Through simulations, we have showed that our proposed pLMMGMM has robust and accurate prediction performance across a range of disease models (Figure 3 and Supplementary Figure S3). In addition, its has also achieved relatively high sensitivity and specificity in correctly detecting prediction regions that harbor genetic variants with various types of predictive effects.

Computational efficiency is one of the major bottleneck for LMMs with multiple random effects (Weissbrod, Geiger and Rosset, 2016; Wen and Lu, 2020; Li, Lu and Wen, 2020). Traditional methods usually obtain the MLE/REML estimators, both of which can be computationally demanding, especially when the number of multiple random effects is large. In our work, we traded computational efficiency with statistical efficiency, and proposed to obtain parameter estimates via the GMM. By using GMM, our objective function is much easier to optimize. Indeed, for REML with traditional Newton-Raphson algorithm, it can barely converge when the number of random effects is above 10. Even for REML with simulated anneal algorithm where the global optimum is not guaranteed (Goffe, Ferrier and Rogers, 1994), the computational time increases at a much faster rate as compared to our proposed method (Figure 2 and Supplementary Figure S2). The computational efficiency allows our proposed method to jointly model a large number of genetic regions and consider various forms of predictive effects, both of which can be important for improving risk prediction models.

In the prediction analyses of FDG and AV45 outcomes, we found that the proposed method outperformed both MKLMM and gBLUP (Figure 4), indicating that the designed variable selection and multi-kernel learning in pLMMGMM can improve the prediction accuracy. In addition, the predictive genes selected by pLMMGMM method are also quite consistent (Supplementary Table S4). The three well-known AD-related genes, *APOC1, APOE* and *TOMM40*, are highly selected (>97%) for both FDG and AV45. The *APOE* gene encodes Apolipoprotein E that is involved in the cholesterol transport (Zannis, Kardassis and Zanni, 1993), and high levels of cholesterol play a significant role in the pathogenesis of AD (Puglielli, Tanzi and Kovacs, 2003). Indeed, *APOE* gene has been identified as a major genetic risk factor for AD in existing literature. The *APOEϵ4* is overrepresented among late-onset AD patients (Strittmatter et al., 1993; Poirier et al., 1993). Tang et al. (1998) found that the presence of *APOEϵ4* is a determinant risk factor of AD in Caucasian, and Graff-Radford et al. (2002) also reported that one or two copies of *APOEϵ4* affect the risk of AD for African American. The *APOC1* gene encodes apolipoprotein C1, a member of apolipoprotein family, and it affects the cholesterol metabolism that is involved in AD pathology (Poirier et al., 1993). In addition, it has also been found that the rs4420638 polymorphism on *APOC1* increases the accumulation of homocysteine, and thus affects AD risk (Prendecki et al., 2018). *TOMM40* regulates the mitochondrial function, and it is also a candidate gene for late-onset AD (Roses, 2010). Roses (2010) found that *rs*10524523 on *TOMM40* is highly associated with late-onset AD. In addition, Prendecki et al. (2018) proposed that *rs*10524523 on *TOMM40* can affect the oxidative damage and thus influence the onset and progression of AD. While our models improved prediction accuracy for both AV45 and FDG, additional replication studies are needed to further investigate the performance of our models.

In summary, we have proposed GMM-based penalized LMMs for risk prediction analyses on high-dimensional genomic data, where the variable selection consistency, variable estimation consistency and asymptotic normality of non-zero parameters have been established. Our proposed pLMMGMM method is highly computationally efficient. It can simultaneously consider a large number of genetic regions, and accurately detect those that are predictive. In addition, our proposed method can accommodate various disease models, as it can select appropriate kernel functions that best reflect the underlying disease model via a data-driven approach.

## Supporting information

Supplementary Figues and Tables

## Acknowledgments

The author(s) wish to acknowledge the use of New Zealand eScience Infrastructure (NeSI) high performance computing facilities, consulting support and/or training services as part of this research. New Zealand’s national facilities are provided by NeSI and funded jointly by NeSI’s collaborator institutions and through the Ministry of Business, Innovation & Employment’s Research Infrastructure programme. URL https://www.nesi.org.nz.

## SUPPLEMENTARY MATERIAL

It contains simulation results for 1000 sample size and disease models description. We also provide the details about genes we used in our paper.

## APPENDIX: PROOFS OF THEOREMS

### A.1. Objective Function

Recall the objective function defined in equation 3 of the main text is:

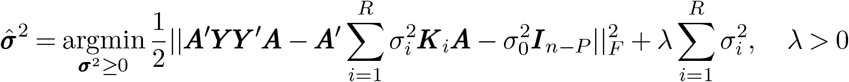

Let 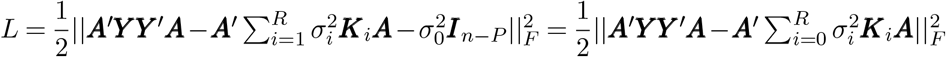 be the loss function, and ***K***_0_ = ***I***_*n–P*_. Then the loss function L can be rewritten as:

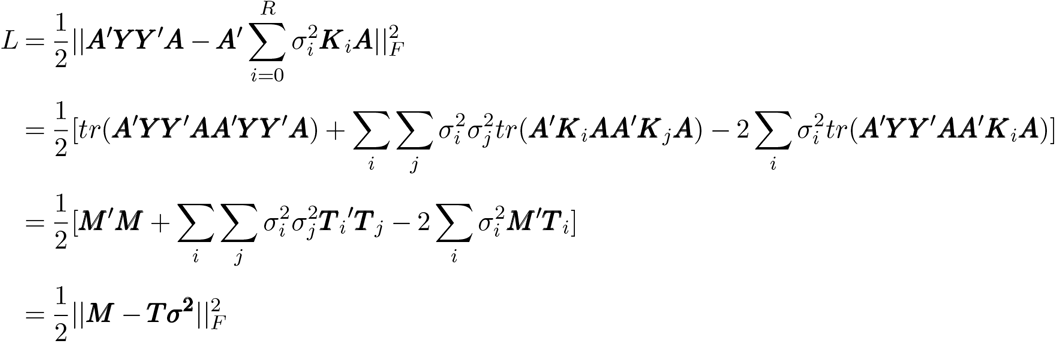

As noted in the main text, ***M*** = *vec*(***A’YY’A***), ***T***_*i*_ = *vec*(***A’K_*i*_A***). Therefore, the estimation of equation 3 in the main text is equal to solving the equation ***M*** = ***Tσ***^2^ + ***ω*** and 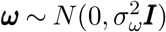. Our penalized objective function can be rewritten as

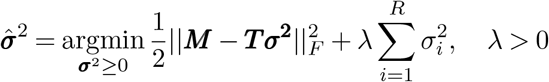

### A.2. Proof of Theorem 2.1

Using the KKT (Karush-Kuhn-Tucker) conditions, 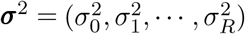 is the non-negative lasso estimator for given λ if and only if:

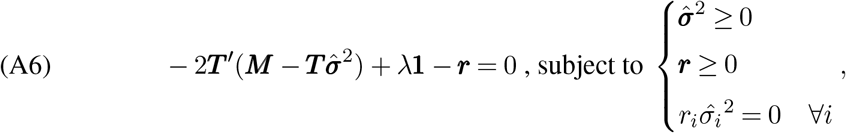

Replace ***M*** by ***Tσ***^2^ + ***ω*** and recall 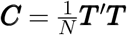. Let 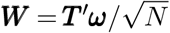. Then the equation A6 can be rewritten as

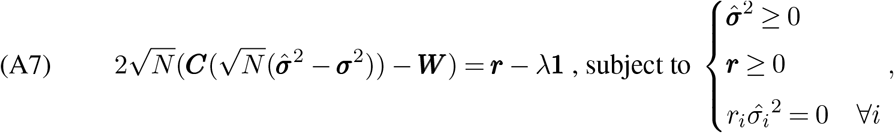

Let 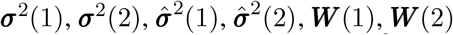 and ***r***(1), ***r***(2) are the *q* + 1 nonzero elements and the *R* – *q* zero elements of 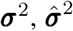, ***W*** and ***r***. If there exist 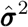 meet equation A7, and 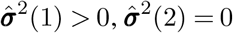. Then *S*_1_ = *S*(λ). That is

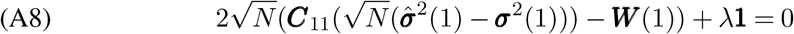

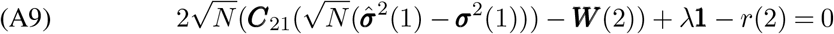

Thus, the existence of 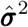 of equation A8 and A9 can be implied by

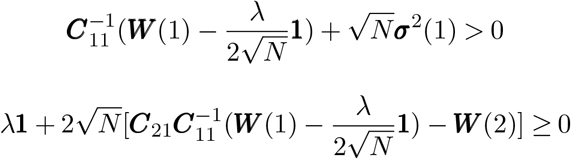

Let 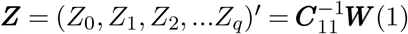 and 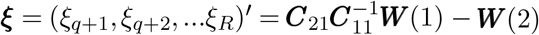, and then

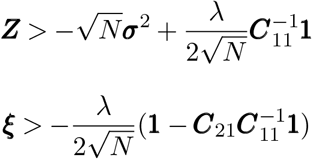

Based on the Nonnegative irrepresentable condition,

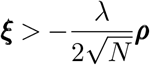

Then the probability is:

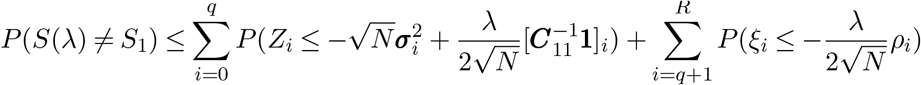

Suppose that 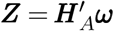 and 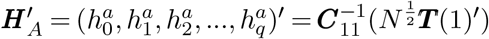. Then

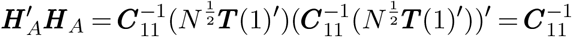

Let ***e***_*i*_ be a vector with *i*th entry one and other zeros.

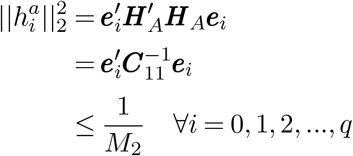

Similarly, suppose that 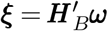 and 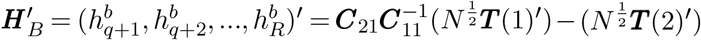. Then

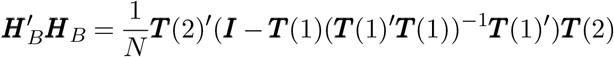

Let ***H*** = ***I*** – ***T***(1)(***T***(1)’***T***(1))^−1^***T***(1)’ and ***H*** is a projection matrix and whose maximal eigen value λ_*max*_(***H***) is 1 or 0.

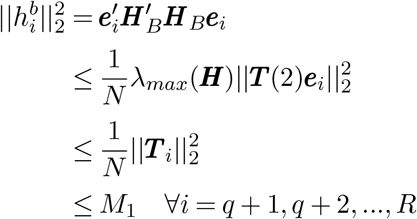

Because ***ω***_*i*_ are i.i.d. Gaussian variables, ***Z***_*i*_ and ***ξ***_*i*_ are also Gaussian variables and are bounded by secondary moment. Let Φ(*t*) denote the distribution function of standard Gaussian variable, then for any *t* > 0,

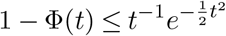

for 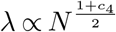, by 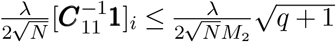, we have

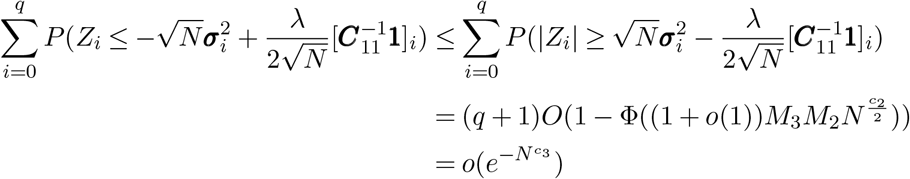

And also we can get

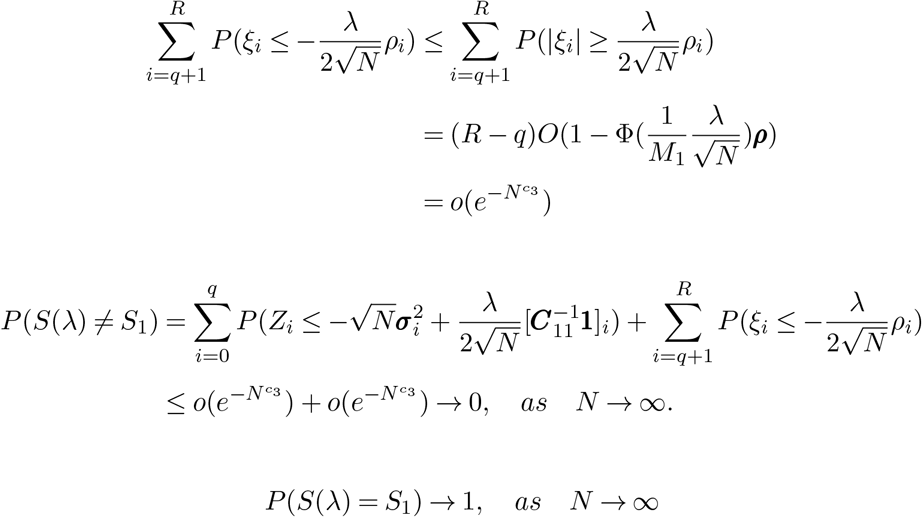

Variable selection consistency proof completed.

### A.3. Proof of Theorem 2.2

Under assumptions 3 and assumptions 4, Negahban et al. (2012) establishes upper bounds for nonnegative lasso estimation errors.

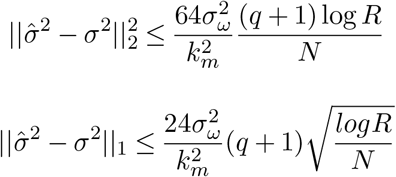

From these bounds, we know that our pLMMGMM method has estimation consistency, i.e., 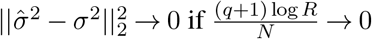, as *N* → ∞

### A.4. Proof of Theorem 2.3

Based on the proof of theorem 1,

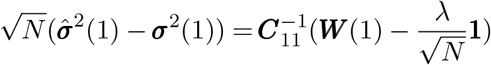

Since 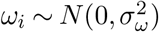, then

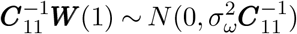

Then we have

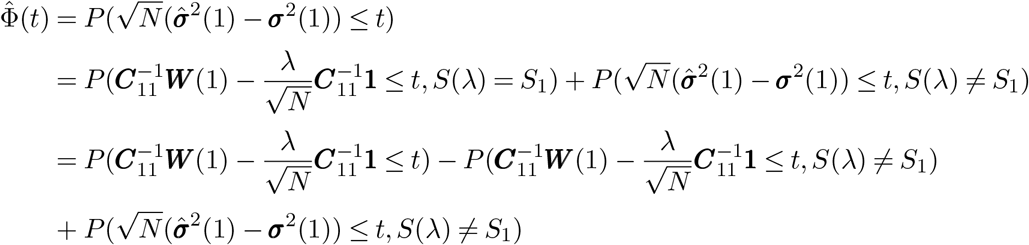

Based on the assumption 1, we can prove that

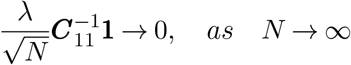

then by Slutsky’s theorem,

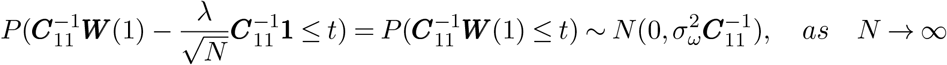

In addition,

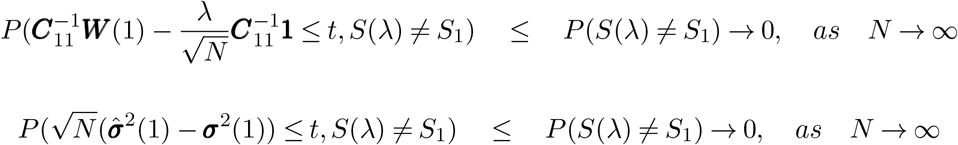

Asymptotic normality proof completed.

