## Supplementary Figues and Tables for "A Penalized Linear Mixed Model with Generalized Method of Moments for Complex Phenotype Prediction"

### Supplementary Materials

Table S1: The chances of selecting two predictive regions as the number of noise regions increases ($n = 1000$).

| Regions | Sensitivity | Specificity |
|---|---|---|
| 5 | 1.000 | 0.894 |
| 10 | 1.000 | 0.887 |
| 50 | 1.000 | 0.905 |
| 100 | 1.000 | 0.922 |

Table S2: Disease models description

| Model | Description | $\boldsymbol{K}_1$ | $\boldsymbol{K}_2$ |
|---|---|---|---|
| $S_1 : L + L$ | Linear additive effects | $k_l(\boldsymbol{x}_1, \boldsymbol{x}_2) =< \boldsymbol{x}_1, \boldsymbol{x}_2 >$ | $k_l(\boldsymbol{x}_1, \boldsymbol{x}_2) =< \boldsymbol{x}_1, \boldsymbol{x}_2 >$ |
| $S_2 : R + R$ | Non-linear effects. | $k_{rbf}(\boldsymbol{x}_1, \boldsymbol{x}_2) = \exp\left[-\frac{1}{2}\|\|\boldsymbol{x}_1 - \boldsymbol{x}_2\|\|_2^2\right]$ | $k_{rbf}(\boldsymbol{x}_1, \boldsymbol{x}_2) = \exp\left[-\frac{1}{2}\|\|\boldsymbol{x}_1 - \boldsymbol{x}_2\|\|_2^2\right]$ |
| $S_3 : P + P$ | Pair-wise interaction effects | $k_p(\boldsymbol{x}_1, \boldsymbol{x}_2) = (< \boldsymbol{x}_1, \boldsymbol{x}_2 >)^2$ | $k_p(\boldsymbol{x}_1, \boldsymbol{x}_2) = (< \boldsymbol{x}_1, \boldsymbol{x}_2 >)^2$ |
| $S_4 : L + R$ | Linear and non-linear effects | $k_l(\boldsymbol{x}_1, \boldsymbol{x}_2) =< \boldsymbol{x}_1, \boldsymbol{x}_2 >$ | $k_{rbf}(\boldsymbol{x}_1, \boldsymbol{x}_2) = \exp\left[-\frac{1}{2}\|\|\boldsymbol{x}_1 - \boldsymbol{x}_2\|\|_2^2\right]$ |
| $S_5 : L + P$ | Linear and pair-wise interaction | $k_l(\boldsymbol{x}_1, \boldsymbol{x}_2) =< \boldsymbol{x}_1, \boldsymbol{x}_2 >$ | $k_p(\boldsymbol{x}_1, \boldsymbol{x}_2) = (< \boldsymbol{x}_1, \boldsymbol{x}_2 >)^2$ |

Table S3: The chances of selecting two predictive regions under different disease models ($n = 1000$).

| Disease models | Sensitivity | Specificity |
|---|---|---|
| $S_1 : L + L$ | 1.000 | 0.911 |
| $S_2 : R + R$ | 1.000 | 0.909 |
| $S_3 : P + P$ | 1.000 | 0.980 |
| $S_4 : L + R$ | 0.999 | 0.904 |
| $S_5 : L + P$ | 0.999 | 0.947 |

Table S4: The chances of selecting genes for real data

| Genes | Chromosome | Start Position | End Position | Probability(FDG) | Probability(AV-45) |
|---|---|---|---|---|---|
| COL11A1 | 1 | 103342022 | 103574052 | 0 | 0 |
| CR1 | 1 | 207669472 | 207815110 | 0 | 0 |
| CR1L | 1 | 207818457 | 207897036 | 0 | 0 |
| FCER1G | 1 | 161185086 | 161189038 | 0 | 0 |
| FLVCR1 | 1 | 213031596 | 213072705 | 0 | 0 |
| FLVCR1-AS1 | 1 | 213029945 | 213031480 | 0 | 0 |
| GBP2 | 1 | 89571815 | 89591842 | 0 | 0 |
| HSD11B1 | 1 | 209859524 | 209908295 | 0 | 0 |
| NGF | 1 | 115828536 | 115880857 | 0 | 0 |
| PARP1 | 1 | 226548391 | 226595801 | 0 | 0 |
| POU2F1 | 1 | 167190065 | 167396582 | 0 | 0 |
| BIN1 | 2 | 127805598 | 127864903 | 0 | 0 |
| LHCGR | 2 | 48913912 | 48982880 | 0 | 0 |
| LRP2 | 2 | 169983618 | 170219122 | 0 | 0 |
| APOD | 3 | 195295572 | 195311076 | 0 | 0 |
| GSK3B | 3 | 119540801 | 119813264 | 0 | 0 |
| SST | 3 | 187386693 | 187388201 | 0 | 0.01 |
| ALB | 4 | 74269971 | 74287129 | 0 | 0.02 |
| COL25A1 | 4 | 109731876 | 110223799 | 0 | 0 |
| ADRB2 | 5 | 148206155 | 148208197 | 0 | 0 |
| ARSB | 5 | 78073036 | 78282357 | 0 | 0 |
| FGF1 | 5 | 141971742 | 142077635 | 0.24 | 0 |
| FGF10 | 5 | 44305096 | 44388784 | 0 | 0 |
| FGF10-AS1 | 5 | 44388833 | 44414091 | 0 | 0 |
| FGF18 | 5 | 170846666 | 170884630 | 0 | 0 |
| NDUFS4 | 5 | 52856464 | 52979171 | 0 | 0 |
| PPP2R2B-IT1 | 5 | 146293769 | 146299069 | 0 | 0 |
| AGER | 6 | 32148744 | 32152099 | 0 | 0.01 |
| HSPA1A | 6 | 31783290 | 31785719 | 0 | 0 |
| MICA | 6 | 31367560 | 31383092 | 0 | 0.01 |
| MICAL1 | 6 | 109765265 | 109787171 | 0 | 0 |
| TBP | 6 | 170863420 | 170881958 | 0 | 0 |
| TBPL1 | 6 | 134273307 | 134308638 | 0 | 0 |
| TREM2 | 6 | 41126243 | 41130924 | 0 | 0 |
| CAV1 | 7 | 116164838 | 116201239 | 0 | 0 |
| PON3 | 7 | 94989183 | 95025687 | 0 | 0.01 |
| RELN | 7 | 103112230 | 103629963 | 0 | 0 |
| ADAM9 | 8 | 38854504 | 38962779 | 0 | 0.03 |
| NAT1 | 8 | 18027970 | 18081198 | 0 | 0 |
| NRG1 | 8 | 31497267 | 32622558 | 0 | 0 |
| DAPK1 | 9 | 90112142 | 90323549 | 0 | 0 |
| DFNB31 | 9 | 117164359 | 117267736 | 0 | 0 |
| HSPA5 | 9 | 127997126 | 128003666 | 0 | 0.01 |
| POMT1 | 9 | 134378288 | 134399193 | 0.05 | 0.04 |
| RXRA | 9 | 137218308 | 137332432 | 0 | 0 |
| TLR4 | 9 | 120466452 | 120479769 | 0 | 0 |
| CACNB2 | 10 | 18429605 | 18830688 | 0 | 0 |
| MINPP1 | 10 | 89264222 | 89313218 | 0 | 0 |
| TET1 | 10 | 70320116 | 70454239 | 0 | 0 |

Table S4: The chances of selecting genes for real data *(continued)*

| Genes | Chromosome | Start Position | End Position | Probability(FDG) | Probability(AV-45) |
|---|---|---|---|---|---|
| TFAM | 10 | 60144902 | 60158990 | 0 | 0 |
| HBG2 | 11 | 5274420 | 5276011 | 0.07 | 0 |
| ATF7 | 12 | 53901639 | 54020199 | 0.03 | 0 |
| ATF7IP | 12 | 14518565 | 14655869 | 0 | 0 |
| OLR1 | 12 | 10310898 | 10324790 | 0 | 0 |
| SLC11A2 | 12 | 51373565 | 51422058 | 0 | 0 |
| KLF5 | 13 | 73629113 | 73651680 | 0 | 0 |
| CINP | 14 | 102814618 | 102829253 | 0 | 0 |
| GNPNAT1 | 14 | 53241910 | 53258386 | 0 | 0 |
| HNRNPC | 14 | 21677295 | 21737638 | 0 | 0 |
| MTHFD1 | 14 | 64854758 | 64926725 | 0.01 | 0 |
| PNP | 14 | 20937537 | 20946165 | 0 | 0 |
| SEL1L | 14 | 81937890 | 82000205 | 0 | 0 |
| SERPINA1 | 14 | 94843083 | 94857029 | 0.21 | 0 |
| SERPINA2 | 14 | 94829974 | 94833039 | 0 | 0 |
| SERPINA3 | 14 | 95078713 | 95090390 | 0 | 0 |
| SERPINA4 | 14 | 95027756 | 95036250 | 0 | 0 |
| SERPINA5 | 14 | 95047705 | 95059457 | 0 | 0 |
| SERPINA6 | 14 | 94770584 | 94789688 | 0 | 0 |
| SERPINA9 | 14 | 94929057 | 94942670 | 0 | 0 |
| SERPINA10 | 14 | 94749649 | 94759608 | 0 | 0 |
| SERPINA11 | 14 | 94908800 | 94919122 | 0 | 0 |
| SERPINA12 | 14 | 94953619 | 94984181 | 0 | 0 |
| SERPINA13P | 14 | 95107061 | 95113331 | 0 | 0 |
| CHRNA3 | 15 | 78885394 | 78913637 | 0 | 0 |
| MEF2A | 15 | 100106132 | 100256629 | 0 | 0 |
| MEFV | 16 | 3292027 | 3306627 | 0 | 0 |
| UBE2I | 16 | 1359153 | 1377019 | 0 | 0 |
| CCL3 | 17 | 34415602 | 34417506 | 0 | 0 |
| CDK5R1 | 17 | 30814104 | 30818271 | 0 | 0 |
| COX10 | 17 | 13972718 | 14111996 | 0 | 0 |
| COX10-AS1 | 17 | 13932608 | 13972775 | 0 | 0 |
| PNMT | 17 | 37824233 | 37826728 | 0 | 0 |
| APOC1 | 19 | 45417920 | 45422606 | 1 | 1 |
| APOE | 19 | 45409038 | 45412650 | 1 | 0.97 |
| GNA11 | 19 | 3094407 | 3124000 | 0 | 0 |
| TOMM40 | 19 | 45394476 | 45406946 | 0.28 | 1 |
| DOPEY2 | 21 | 37536838 | 37666572 | 0 | 0 |
| MCM3AP | 21 | 47655038 | 47705308 | 0 | 0 |
| MCM3AP-AS1 | 21 | 47649144 | 47671615 | 0 | 0 |
| NCAM2 | 21 | 22370632 | 22912517 | 0 | 0 |
| RUNX1-IT1 | 21 | 36410232 | 36411723 | 0.07 | 0 |
| S100B | 21 | 48018530 | 48025035 | 0.24 | 0 |
| SAMSN1 | 21 | 15857548 | 15955723 | 0 | 0 |
| SAMSN1-AS1 | 21 | 15954522 | 15970624 | 0.01 | 0 |
| SEPT3 | 22 | 42372930 | 42394225 | 0 | 0 |

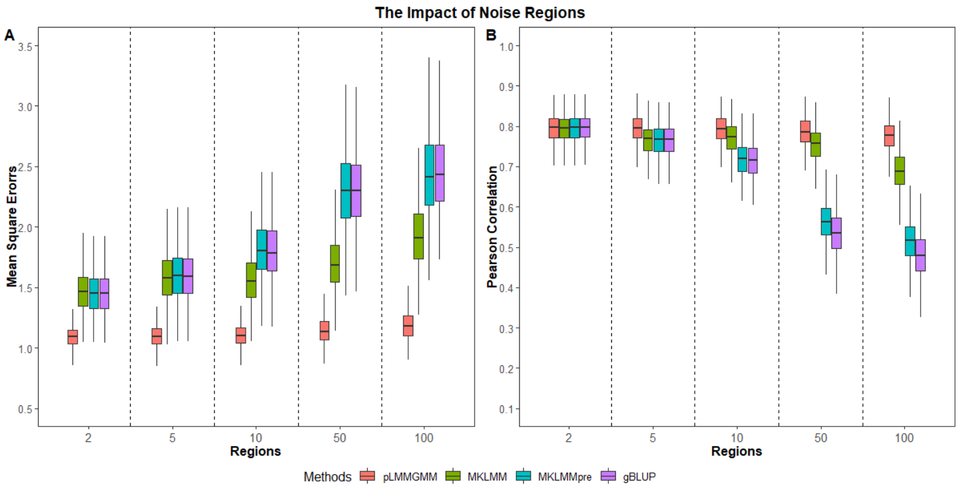


Figure S1: The impact of the number of noise regions on Pearson correlations and MSEs ($n = 1000$).

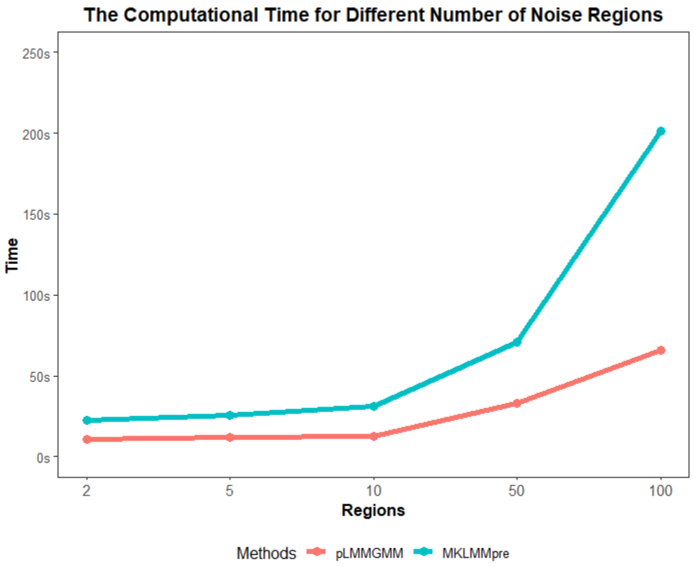


Figure S2: The impact of the number of noise genomic regions on computational time ($n = 1000$).

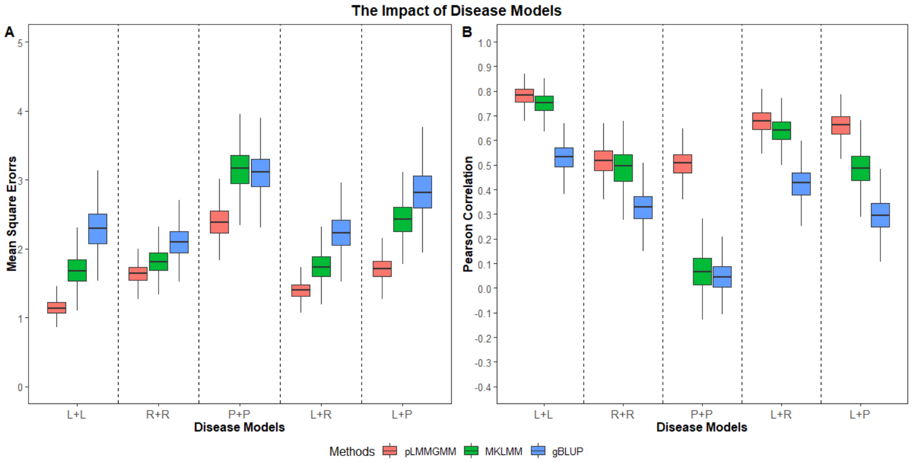


Figure S3: The impact of disease models. **L+L:** genetic variants on both regions have linear additive effects. **R+R:** predictors from both regions have non-linear predictive effects. **P+P:** both regions harbor variants with pair-wise interaction effects. **L+R:** genetic variants on the first and second regions have linear additive and non-linear effects, respectively. **L+P:** predictors on the first and second regions have linear additive and pair-wise interaction effects, respectively ($n = 1000$).

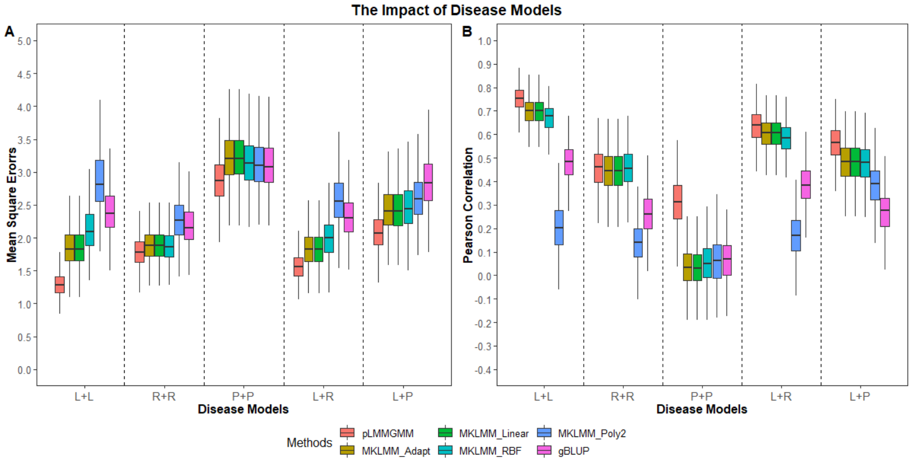


Figure S4: The impact of disease models. **L+L:** genetic variants on both regions have linear additive effects. **R+R:** predictors from both regions have non-linear predictive effects. **P+P:** both regions harbor variants with pair-wise interaction effects. **L+R:** genetic variants on the first and second regions have linear additive and non-linear effects, respectively. **L+P:** predictors on the first and second regions have linear additive and pair-wise interaction effects, respectively ($n = 500$).

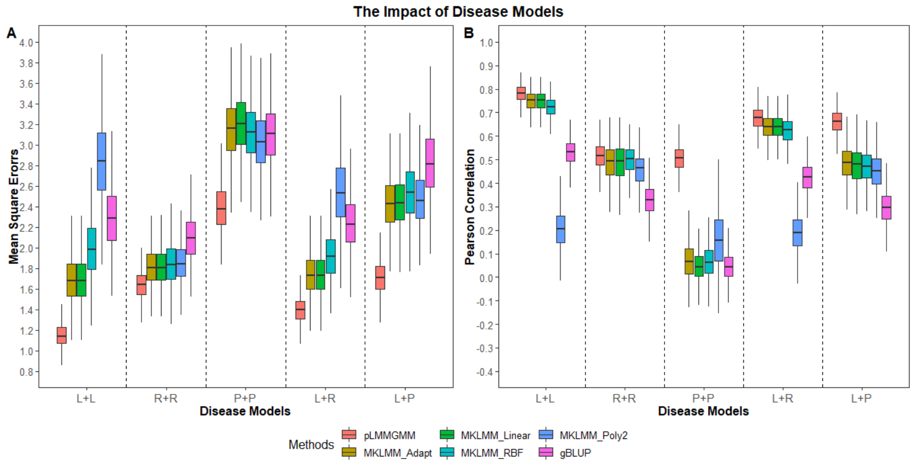


Figure S5: The impact of disease models. **L+L:** genetic variants on both regions have linear additive effects. **R+R:** predictors from both regions have non-linear predictive effects. **P+P:** both regions harbor variants with pair-wise interaction effects. **L+R:** genetic variants on the first and second regions have linear additive and non-linear effects, respectively. **L+P:** predictors on the first and second regions have linear additive and pair-wise interaction effects, respectively ($n = 1000$).

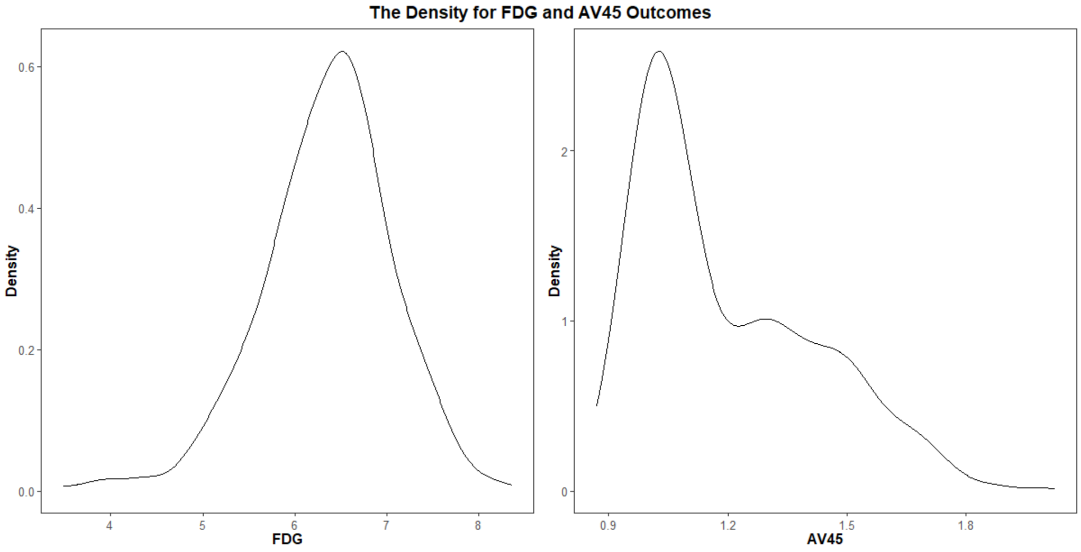


Figure S6: The distribution for FDG and AV45 outcomes.
